# A nuclear hormone receptor *nhr-76* programs age-dependent chemotaxis decline in *C. elegans*

**DOI:** 10.1101/2024.08.30.609799

**Authors:** Rikuou Yokosawa, Kentaro Noma

## Abstract

A decline in food-searching behavior of post-reproductive animals can be beneficial for the population and possibly programmed by the genome. We investigated the genetic program of age-dependent decline in chemotaxis behavior toward an odorant secreted from bacterial food in *C. elegans*. Through a forward genetic screen, we identified a nuclear hormone receptor, *nhr-76*, whose mutants ameliorate the age-dependent chemotaxis decline. We found that *nhr-76* downregulates the expression of the odorant receptor during aging. Because NHR-76 expression and localization did not change during aging, secretion of its hydrophobic ligands might alter the activity of NHR-76 to cause age-dependent chemotaxis decline. Our findings imply that post-reproductive behavioral decline can be genetically programmed.

## Introduction

Aging can be defined as the functional decline of an organism after sexual maturation. Passive accumulation of cellular and molecular damages such as cellular senescence, genomic instability, and loss of proteostasis are widely accepted to cause aging (*1*). In contrast, the active genetic program that causes aging, especially in naturally aged animals, has been less explored. Among aging phenotypes, behaviors can impact the fitness of a population. Thus, we speculate that behavioral aging can be genetically programmed for a population even after the end of reproduction. *C. elegans* is ideal for studying behavioral aging due to its short lifespan and robust behavior with a simple nervous system. Chemotaxis behavior is an innate ability to sense and migrate toward volatile odorants (Fig. 1A) (*2*). For example, *C. elegans* chemotax toward diacetyl, derived from bacterial food (*3*). Diacetyl is sensed by the ODR-10 receptor specifically expressed in the AWA sensory neurons (*2, 4*). We used the chemotaxis behavior toward diacetyl to examine the genetic program of behavioral aging.

**Figure 1.**
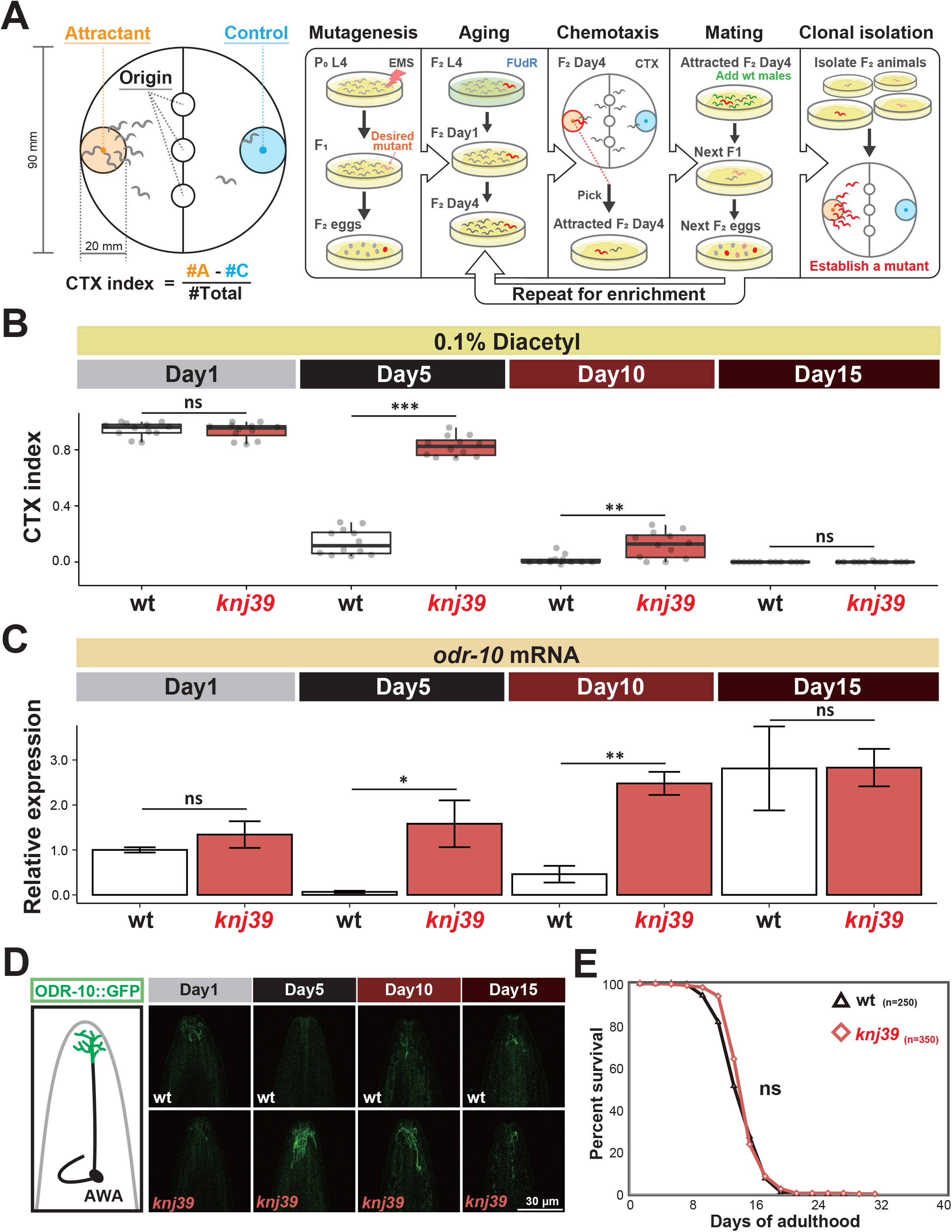
Forward genetic screen identified a mutant that ameliorated age-dependent chemotaxis decline. (**A**) Schematic of population chemotaxis assay and the forward genetic screen to isolate mutants ameliorating age-dependent chemotaxis decline. Animals’ ability chemotax toward an attractive odorant was examined. The chemotaxis (CTX) index was calculated using the indicated formula, where #A and #C indicate the number of animals in the attractant and control circles, respectively. The steps from Aging to Mating were repeated multiple times to enrich the desirable mutants. (**B**) *knj39* mutants ameliorated age-dependent chemotaxis decline toward 0.1% diacetyl on Day5. (**C**) The *odr-10* mRNA expression normalized by the mean value of the Day1 wild type. Error bars indicate SEM. (**D**) Schematic of the head of an animal and representative images of the ODR-10::GFP reporter, expressed in AWA neurons. Scale bar=30 µm. (**E**) *knj39* mutants did not live longer than wild-type animals. n indicates the number of animals examined. Statistical tests were conducted using Mann-Whitney U test between same-day conditions for (**B**), Student’s t-test between same-day conditions for (**C**), and the Log-rank test for (**E**). ns: p>0.05; *p<0.05; **p<0.01.

## Results

### Forward genetic screen identified the *knj39* mutant that ameliorated age-dependent chemotaxis decline

As we reported previously, the chemotaxis ability toward diacetyl and the transcripts and GFP-tagged protein expression of the diacetyl receptor, *odr-10*, declined from young (Day1 of adulthood, Day1) to aged adults (Day5) after the end of self-reproduction of a hermaphrodite (*5*) (Fig. 1B,1C,1D wt). Further analysis revealed that the expression of *odr-10* was increased from Day5 to Day10 when most animals were still alive, although the chemotaxis ability remained low on Day10 (Fig. 1B,1C,1D, wt). We speculated that an active genetic mechanism might actively decrease the *odr-10* expression and the chemotaxis ability of Day5 animals.

To isolate mutants regulating age-dependent chemotaxis decline, we sought to perform a forward genetic screening. However, the mass screening with Day5 hermaphrodites was not feasible because they did not have self-fertilized progeny (fig. S1, without males or FUdR). Since sperm depletion terminates self-reproduction (*6*), we mated aged hermaphrodites with young males and could obtain progeny even from aged animals (fig. S1, Materials and Methods). Thus, we performed a forward genetic screen based on the chemotaxis ability of the F_2_ progeny of mutagenized animals on Day4 (Fig. 1A). A few rounds of mating and behavioral assays eliminated false positives and enriched potential mutants (Fig. 1A, Materials and Methods).

From this screen, we isolated a mutant *knj39* that ameliorated the chemotaxis decline toward 0.1% diacetyl on Day5 (Fig. 1B, *knj39*). Consistent with the chemotaxis behavior, the expression of *odr-10* transcripts and GFP-tagged ODR-10 did not decline from Day1 to Day5 in the *knj39* mutants (Fig. 1C,1D, *knj39*). Day15 *knj39* mutants showed low chemotaxis indices like the wild type, while the *odr-10* expression was high (fig. 1B,1C, *knj39*). This low chemotaxis ability might be due to locomotion defects (fig. S2) and/or defective integration of sensory information rather than lower sensory function. The chemotaxis index of *knj39* heterozygotes was between the wild type and *knj39* homozygotes, suggesting that *knj39* was semi-dominant (fig. S3). The better chemotaxis of aged *knj39* mutants was not due to the secondary effects of the better chemotaxis ability of young animals (fig. S4) or longer lifespan (Fig. 1E). Although the timing of the chemotaxis decline is correlated with the end of the self-reproduction (*5*), the self-reproductive span of *knj39* was comparable to the wild type (fig. S5 A, B).

Like Day1 wild-type animals (*4*), Day5 *knj39* mutants required *odr-10* for the chemotaxis behavior (fig. S6), suggesting that aged *knj39* mutants did not recruit other receptors to achieve chemotaxis toward diacetyl. Collectively, these results suggest that *knj39* mutants lack a genetic program to decrease *odr-10* expression and chemotaxis behavior in aged animals.

### NHR-76 functions in AWA neurons to cause age-dependent chemotaxis decline

Based on the chromosome linkage analysis and whole-genome sequencing, *knj39* was predicted to be a mutation of *nhr-76* encoding a nuclear hormone receptor (Fig. 2A). *knj39* had a substitution mutation from an arginine to a stop codon in *nhr-76* (Fig. 2A). In the chemotaxis assays, three additional deletion mutants of *nhr-76* (*tm671, knj51*, and *knj52*) phenocopied *knj39* mutants (Fig. 2B); the genomic *nhr-76* fragment rescued *nhr-76(knj51*) (Fig. 2C, fig. S7, *nhr-76*p). These results suggest that the loss of function of *nhr-76* ameliorated age-dependent chemotaxis decline. A previous study showed that *nhr-76* regulates lipid metabolism in the intestine (*7*). However, the intestine-specific expression did not rescue the *nhr-76(knj51*) deletion mutants in the age-dependent chemotaxis decline (Fig. 2C, fig. S8). Moreover, the mutants in the *nhr-76*-related lipid metabolism pathway (*7*) showed no phenotype of age-dependent chemotaxis decline (fig. S8). These results suggest that *nhr-76* ameliorates the age-dependent chemotaxis decline independently from the regulation of lipid metabolism. In contrast, the expression of *nhr-76* in all neurons or only in AWA sensory neurons rescued *nhr-76(knj51*) deletion mutants. Thus, *nhr-76* regulates age-dependent chemotaxis decline in AWA, where the diacetyl receptor *odr-10* is specifically expressed (*4*).

**Figure 2.**
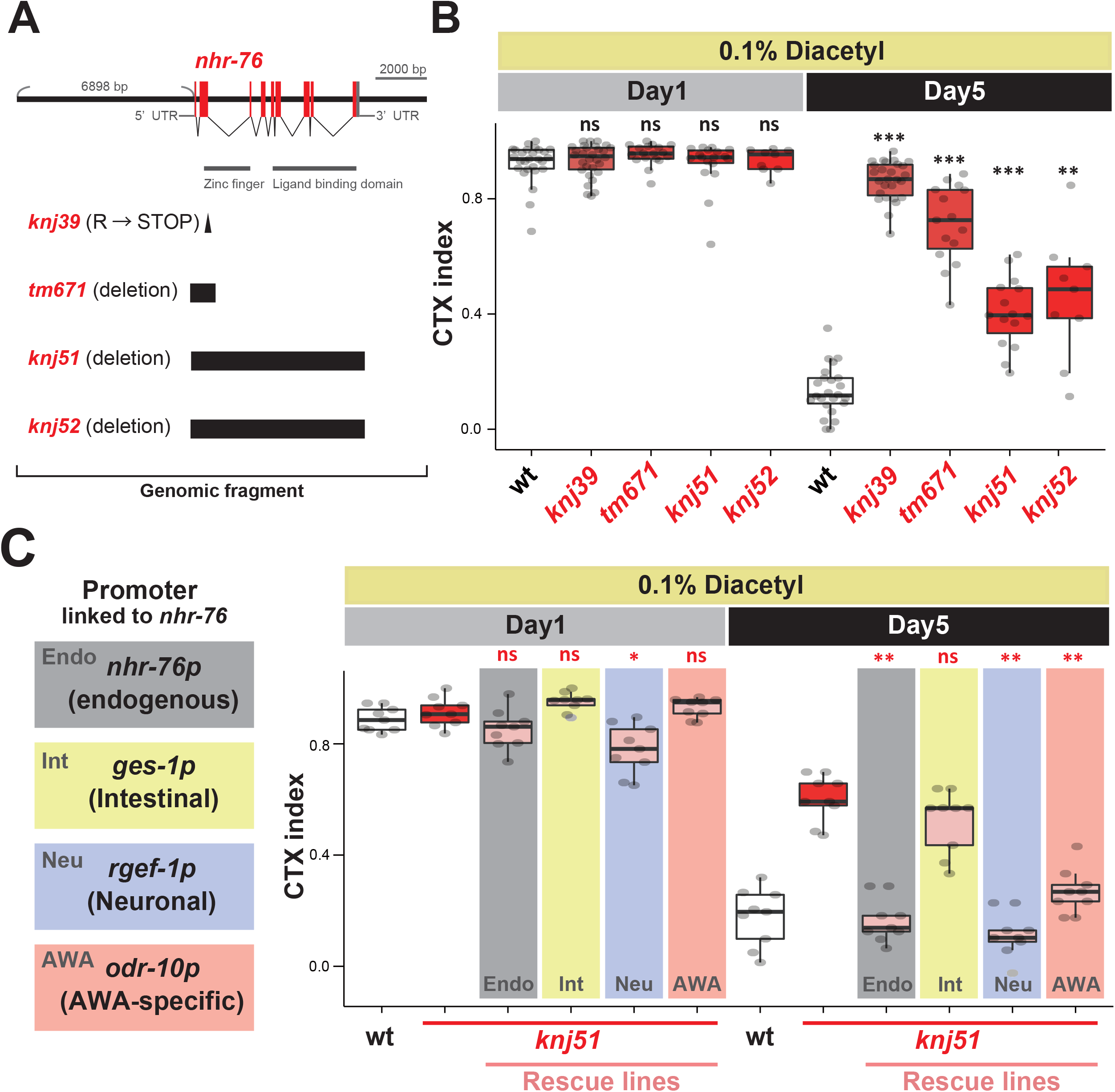
*nhr-76* functions in AWA neurons. (**A**) Gene structure and alleles of *nhr-76*. Red boxes, exon; triangle, point mutation; black box, deletion. (**B**) Like *knj39*, deletion mutants of *nhr-76* ameliorated chemotaxis toward 0.1% diacetyl on Day5. (**C**) The expression of the genomic fragment of *nhr-76* and the pan-neuronal and AWA-specific expression of *nhr-76* cDNA rescued age-dependent chemotaxis decline, but the intestine-specific expression did not. Statistical tests were conducted using Kruskal-Wallis with Steel test against the wt and *knj51* mutant for (**B)** and (**C**), respectively. ns: p>0.05; *p<0.05; **p<0.01; ***p<0.001.

### NHR-76 may change its activity age-dependently to inhibit ODR-7

In young animals, *odr-7* encoding a nuclear hormone receptor is required for chemotaxis toward diacetyl by promoting the *odr-10* expression (*8, 9*). To investigate how *nhr-76* regulates the *odr-10* expression, we analyzed the epistasis between *nhr-76* and *odr-7*. Like young wild-type animals, *odr-7* was required for chemotaxis behavior (Fig. 3A) and the *odr-10* expression (Fig. 3B) of the aged *nhr-76* mutants. Thus, *nhr-76* regulates *odr-10* expression through *odr-7*. Since the expression of *odr-7* did not drastically change during aging (Fig. 3C) (*5*), *nhr-76* may inhibit the activity, but not the expression, of *odr-7* to decrease the *odr-10* expression. *odr-7* is also required for chemotaxis toward pyrazine sensed by a different receptor from *odr-10* in AWA sensory neurons (*9*). In contrast to diacetyl chemotaxis, *nhr-76* mutants did not ameliorate the age-dependent chemotaxis decline toward pyrazine (Fig. 3D) and toward benzaldehyde sensed by another pair of sensory neurons, AWC (*2*) (fig. S9). These results suggest that NHR-76 specifically inhibits the ODR-7 activity for the *odr-10* expression.

**Figure 3.**
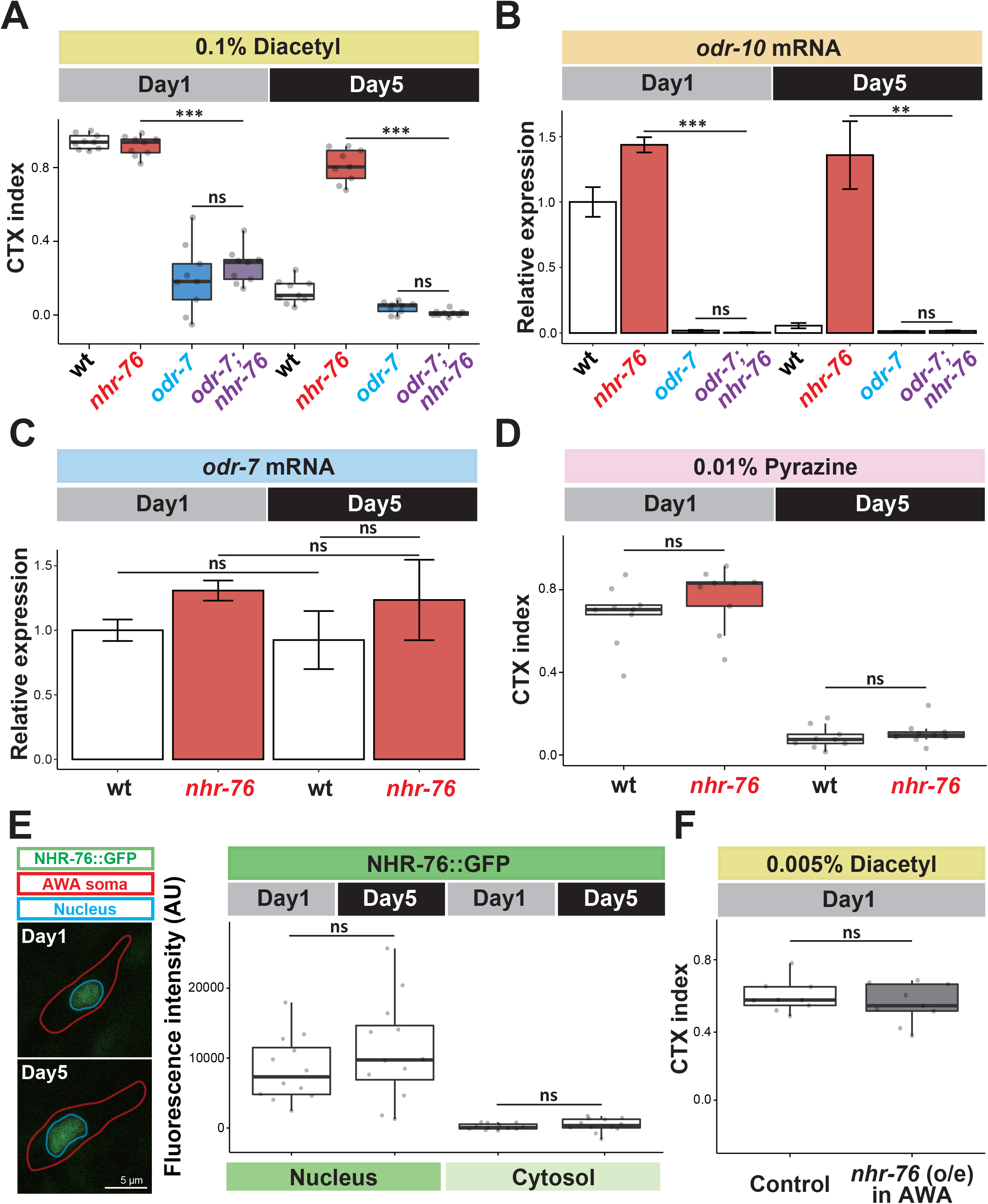
*nhr-76* regulates chemotaxis toward diacetyl through *odr-7*. (**A**) *nhr-76 (knj39*) mutants required *odr-7* for chemotaxis toward diacetyl on Day1 and Day5. (**B**) The mRNA expression of *odr-10* was *odr-7*-dependent in the wild-type and *nhr-76(knj39*). (**C**) *odr-7* mRNA expression did not change between Day1 and Day5 in the wild-type and *nhr-76(knj39*). (**D**) Wild-type animals and *nhr-76(knj39*) mutants showed age-dependent chemotaxis decline toward 0.01% pyrazine. **(E)** The fluorescence intensity of NHR-76::GFP reporter in AWA sensory neurons did not change between Day1 and Day5. AWA sensory neuron was identified with *odr-10p::tagRFP*. The red and blue lines indicate the AWA cell body and the nucleus, respectively. **(F)** *nhr-76* overexpression in AWA sensory neurons did not decrease chemotaxis ability toward 0.005% diacetyl on Day1. Statistical tests were conducted using Kruskal-Wallis with Steel tests against *odr-7; nhr-76* double mutant for (**A**), One-way analysis of variance (ANOVA) with Dunnett’s test against *odr-7; nhr-76* double mutant for (**B**), One-way ANOVA with Tukey’s test for (**C**), Kruskal-Wallis with Steel-Dwass test for (**D**), Student’s t-test for (**E**), and Mann-Whitney U test for (**F**). ns: p>0.05; *p<0.05; **p<0.01.

To address how NHR-76 can inhibit ODR-7 age-dependently, we observed the NHR-76::GFP expression in AWA neurons. The expression and nuclear localization of NHR-76 did not change from Day1 to Day5 (Fig. 3E). Moreover, the overexpression of *nhr-76* in AWA neurons did not decrease chemotaxis ability toward diacetyl in young animals (Fig. 3F, fig. S10). These results suggest that NHR-76 might require a ligand specifically expressed in aged animals to inhibit ODR-7.

### NHR-76 regulates the age-dependent decline of chemotaxis toward diacetyl-producing bacteria

What is the physiological relevance of chemotaxis decline toward diacetyl? Diacetyl is an odor of bacterial food for *C. elegans*, and *Lactobacillus paracasei* (*L. paracasei*) cultured in MRS media supplemented with glucose and pyruvate produces diacetyl (Fig. 4A) (*10*). Young *C. elegans* were attracted toward an agar plug with *L. paracasei* placed on the lid of an assay plate (Fig. 4B,4C, Day1, wt) (*3*). Like the diacetyl chemotaxis, chemotaxis toward *L. paracasei* was reduced in the aged wild type, while aged *nhr-76* mutants maintained the ability to reach *L. paracasei* (Fig. 4C). These results suggest that chemotaxis decline toward diacetyl caused by *nhr-76* possibly impacts the food-searching behavior of aged animals.

**Figure 4.**
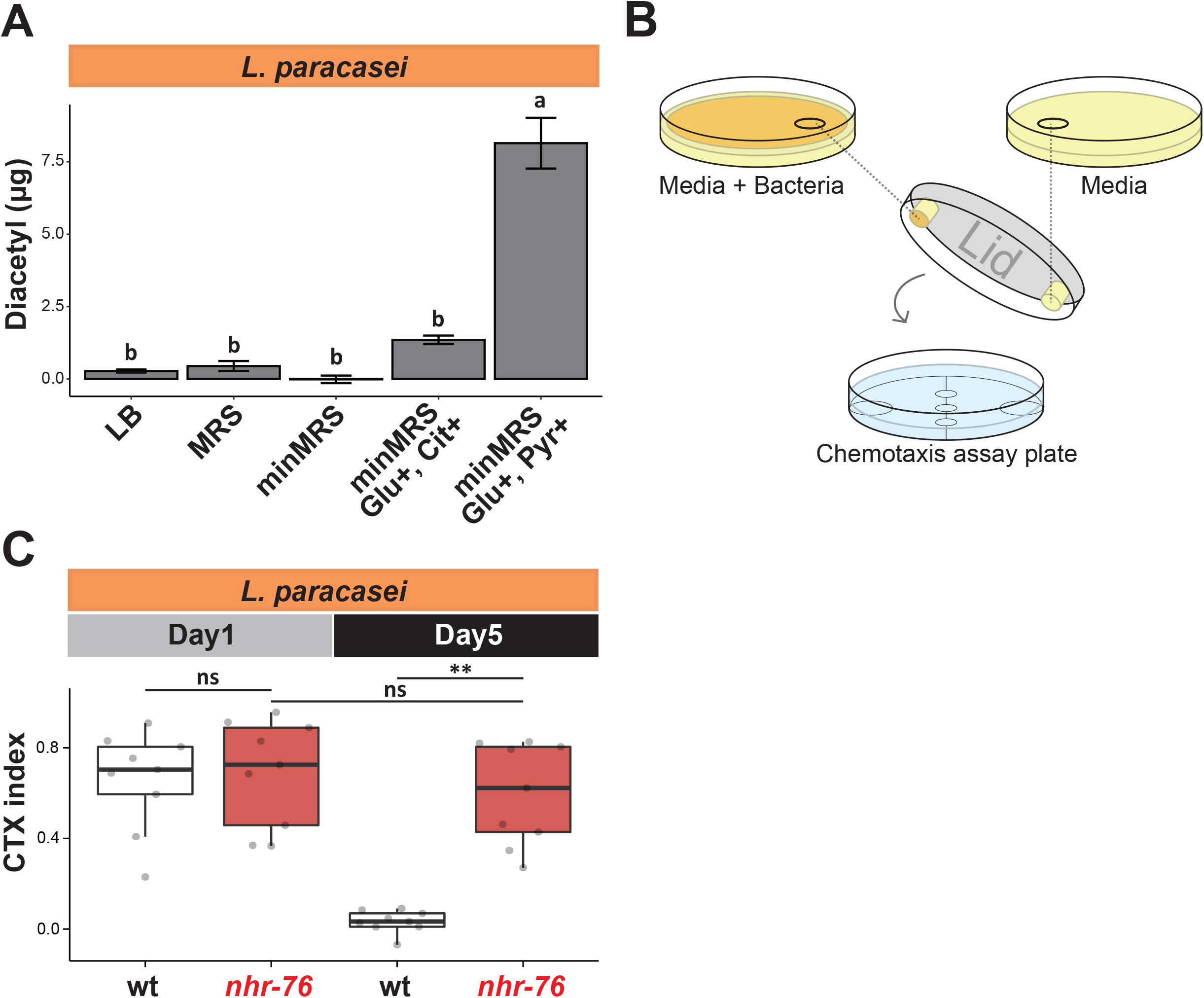
*nhr-76* mutants ameliorate age-dependent chemotaxis decline toward diacetyl-producing bacteria. (**A**) *L. paracasei* produced diacetyl on minimal MRS agar plates supplemented with glucose and pyruvate. (**B**) Schematic of food-taxis assay. The agar plugs with or without *L. paracasei* were placed on the lid of assay plates. (**C**) *nhr-76(knj39*) mutants, but not the wild type, maintained food-taxis ability toward odor of *L. paracasei* on Day5. Statistical tests were conducted using One-way ANOVA with Tukey’s test for (**A**) and Kruskal-Wallis with Steel-Dwass test for **(C**). Different alphabets indicate significant differences. ns: p>0.05; **p<0.01.

**Figure 5.**
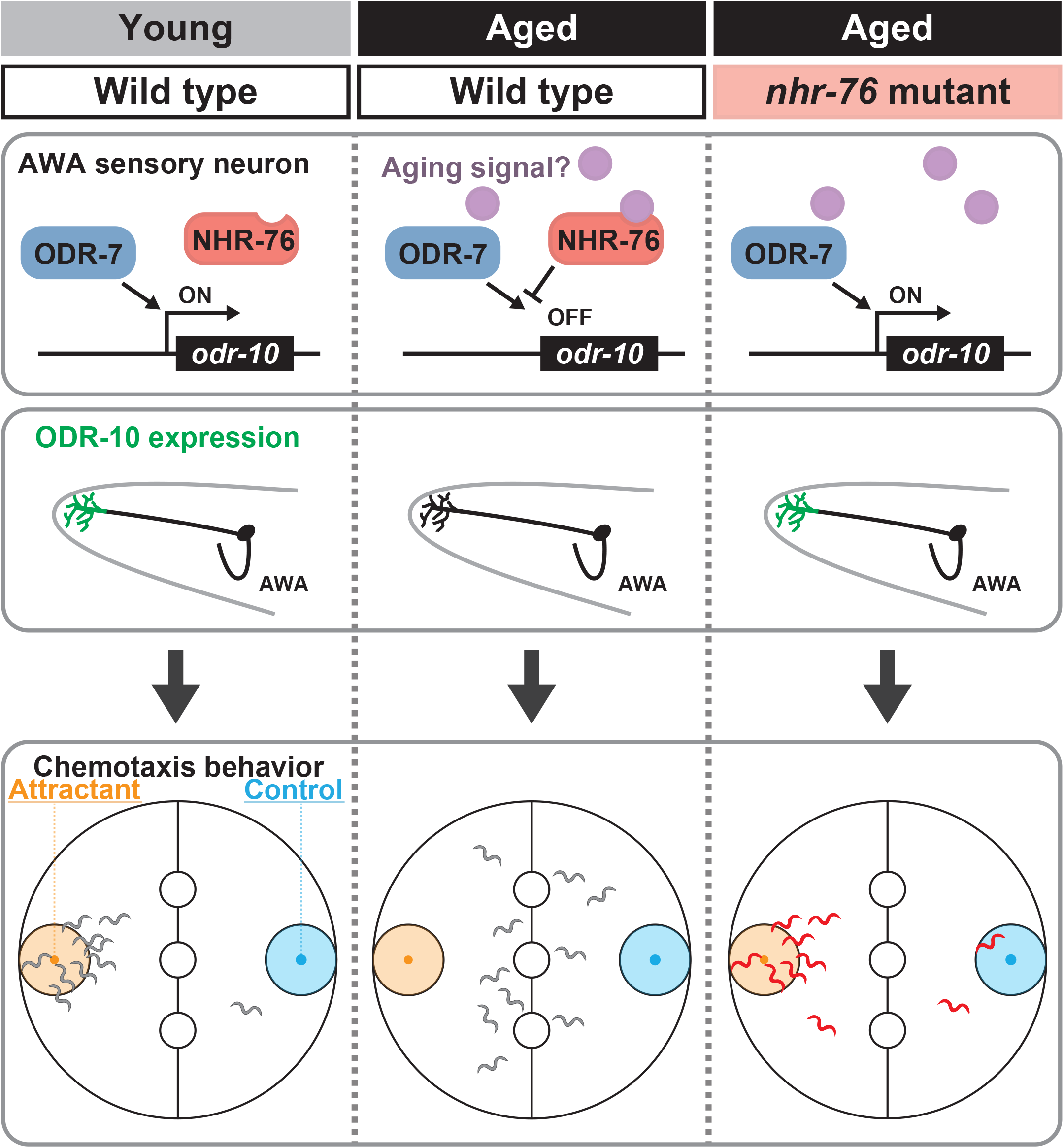
Model of age-dependent chemotaxis decline. *odr-10* encoding diacetyl receptor is expressed through ODR-7 in young animals. In aged animals, NHR-76 is activated by unknown signals shown in purple and inhibits *odr-10* mRNA expression, resulting in the decline of chemotaxis behavior toward diacetyl. In *nhr-76* mutants, the lack of *odr-10* inhibition maintains the ODR-10 expression and the chemotaxis behavior in aged animals.

## Discussion

Forward genetic screens for age-associated phenotypes were limited even in *C. elegans* because aged animals with a phenotype of interest would have few or no progeny. As a result, its application has been only reported for early-onset defects like the progressive deficit in locomotion (*11*) or increased protein aggregation (*12*). Here, we demonstrated that forward genetic screening using aged animals is possible by mating aged hermaphrodites with young males. This novel screening strategy enabled us to find the *nhr-76* mutant, which ameliorates an age-dependent behavioral defect. *nhr-76* had a specific age-dependent role in AWA neurons different from the function in the intestine of young animals (*7*). Moreover, the *nhr-76* mutation did not sacrifice the fitness, based on the brood size (fig. S5 C, D). Thus, *nhr-76*’s function in aging cannot be explained by an antagonistic pleiotropy (*13, 14*). Instead, *nhr-76* appears to actively cause age-dependent chemotaxis decline. In the future, forward genetic screening can be applied to various age-associated phenotypes manifested at relatively earlier stages (*15-17*) to reveal active mechanisms of aging.

Two nuclear hormone receptors, ODR-7 and NHR-76, function as positive and negative regulators of *odr-10* expression. Since *odr-7* affects both diacetyl and pyrazine and *nhr-76* only affects diacetyl, NHR-76 may specifically recognize the *odr-10* promoter region instead of inhibiting the entire ODR-7 function. Since the expression and nuclear localization of NHR-76 did not change in aged animals, we speculate that NHR-76 gets activated during aging. Nuclear hormone receptors can bind to their ligands, such as steroid hormones and hydrophobic vitamins, to change the expression of the target genes (*18*). A hydrophobic ligand may function as a systemic “aging signal” to activate NHR-76. NHR-76 may link the aging signal to a specific behavior. This hypothesis predicts that other nuclear hormones cause behavioral declines of chemotaxis toward other odorants and other food-searching behaviors. Indeed, a nuclear hormone receptor *nhr-66* causes the age-dependent decline in long-term associative memory in *C. elegans (19*).

We showed that *C. elegans* are innately attracted by a diacetyl-producing lactic acid bacterium, *L. paracasei*. Although *L. paracasei* did not support the growth of *C. elegans* (fig. S11), *C. elegans* in a natural habitat should be attracted to the mixture of bacteria as a food source, including lactic acid bacteria (*20*). Since the chemotaxis decline is coupled with the end of reproduction (*5*), post-reproductive individuals might be actively programmed to decrease their attraction to a food source and avoid competing with young animals or mating with them. The active mechanism might also influence associative learning behavior for food because we found that aberrant activation of neurons interferes with the intact circuit to cause age-dependent behavioral decline (*21*). Since age-dependent changes in the nervous system can directly affect the population of a species, an active genetic program might be evolved to couple the end of reproduction with behavioral decline to increase the fitness of a species.

## Materials and Methods

### *C. elegans* culture and strains

Bristol N2 hermaphrodites were used as the wild-type *C. elegans. C. elegans* were cultured at 23°C and fed with *E. coli* OP50 strain on Nematode Growth Medium (NGM) agar plates (*22*). OP50 was precultured in LB broth for 16 hours at 37°C, and 250 µl were seeded on NGM agar plates. Age-matched animals were obtained by preparing synchronized eggs with a bleaching solution (1:1 mixture of 1 M NaOH and household bleach) (*23*). Unless otherwise noted, animals were treated with 25 µM 2’-Deoxy-5-fluorouridine (FUdR) 48 hours after preparing eggs to prevent progeny from hatching. The animals grown at 23°C for 3 days, 6 days, and 7 days from eggs were considered Day1, Day4, and Day5 adults, respectively. We used survivors for Day10 and Day15 when some animals started dying (Fig. 1D). The mutants and transgenic strains used in this work are summarized in Table S1. Transgenic animals carrying extrachromosomal arrays were selected by 0.125 mg/ml hygromycin in NGM plates from eggs for behavioral assays.

### Population chemotaxis assay

Population chemotaxis assays were conducted at 23°C as previously described (*2, 5*). Age-synchronized animals were collected with M9 buffer (20 mM KH_2_PO_4_, 20 mM Na_2_HPO_4_, 8 mM NaCl_2_, and 20 mM NH_4_Cl) and washed with M9 buffer once and CTX buffer (5 mM K-PO_4_, 1 mM CaCl_2_, and 1 mM MgSO_4_) three times. The animals were spotted on the three center circles of a 90-mm chemotaxis plate containing 12 ml agar (2% Difco agar, 1 mM MgSO_4_, 1 mM CaCl_2_, 5 mM K-PO_4_) (Fig. 1A). Along with 1 µl of NaN_3,_ 4 µl of an attractant diluted with 100% ethanol and 4 µl ethanol were placed on the centers of the attractant and control circles, respectively (Fig. 1A). The assay plates were sealed with parafilm and kept in the dark for 1 hour. The animals were killed with chloroform after the assay, and the number of animals in each section was counted under a dissection microscope. CTX index was calculated using the formula indicated in Fig. 1A.

### Locomotion assay

Locomotion speed was measured as the migration distance on a bacterial food lawn at 23°C. A single animal was transferred onto the OP50 lawn of a new NGM plate by picking and removed by picking 1.5 minutes after transferring. The image of the track on the OP50 lawn was taken with a camera (Panasonic HC-V620M-S) and a microscope adapter (MICRONET NY-VS 811383) under a dissection microscope. The length of a track was measured using ImageJ (*24*).

### Forward genetic screen

P_0_ L4 animals were mutagenized with 47 mM EMS for 4 hours (*22*). F_1_ gravid adults were treated with a bleaching solution to obtain synchronized F_2_ eggs. Age-synchronized F_2_ animals were treated with 25 µM FUdR 48 hours after preparing eggs. Twenty-four hours after FUdR treatment, F_2_ animals were washed off with nematode growth (NG) buffer (51 mM NaCl, 1 mM CaCl_2_, 1 mM MgSO_4_, 25 mM K-PO_4_) and transferred onto new NGM plates with OP50 to remove FUdR and mutants with developmental defects. The chemotaxis assay toward 0.01% diacetyl was conducted with Day4 F_2_ animals (6 days after preparing eggs) for 30 minutes without NaN_3_. The attracted Day4 F_2_ animals were pooled, transferred to new NGM plates, and mated with young wild-type males. These animals were considered as P_0_ of the next round of the screening, and the process of chemotaxis and mating was repeated to reduce false positives and enrich mutants until the CTX index became significantly high (Fig. 1A). After this cycle, 8 to 16 F_2_ animals were singly isolated, and the chemotaxis ability was tested on Day4 and Day5. The mutant consistently showing a high CTX index was named *knj39*.

### Quantitative RT-PCR

Age-synchronized animals were used for RNA extraction and cDNA synthesis. Animals grown at 23°C for 56 hours (non-gravid adults), instead of 72 hours for chemotaxis assay, were considered Day1 to exclude the contribution of eggs. RNAiso plus (Takara Bio) and ReverTra Ace® (TOYOBO) were used for RNA extraction and cDNA synthesis, respectively. Quantitative polymerase chain reaction (qPCR) was performed with THUNDERBIRD® SYBR™ qPCR Mix (TOYOBO) using LightCycler®96 (Roche) and gene-specific primers described in Table S2. *cdc-42*, showing a stable expression during aging, was used as a reference (*25*). The mean value of three technical replicates was considered as one biological replicate, and 3 or more biological replicates were examined for one condition.

### Confocal microscopy

Animals were immobilized in a drop of M9 containing 5 mM levamisole on a 4% agarose pad. Images were taken by a confocal microscope LSM880 (Zeiss) with 488-nm and 561-nm lasers for GFP and tagRFP, respectively. The positions of AWA sensory neurons were determined by *odr-10*p::tagRFP. The GFP fluorescence intensities of the nucleus and cytosol were quantified using ImageJ (*24*).

### Lifespan assay

Animals were treated with 25 µM FUdR starting from 48 hours after preparing eggs. Twenty-five animals were transferred to an individual plate on Day1. The animals that did not respond to touch were considered dead and counted every other day; the animals that escaped from the plates were considered censored. After scoring, all living animals were transferred to a new NGM plate containing 25 µM FUdR by picking. The analyses were conducted with OASIS2 (online application for survival analysis 2) (*26*).

### Plasmids

Gateway system (Thermo Fisher Scientific) was used to generate expression plasmids from the entry clone and destination plasmids using LR reaction. Entry clones and destination plasmids were generated by Gibson assembly (*27*). Plasmids were summarized in Table S3.

### CRISPR KO and transgenic strains

CRISPR KO strains were generated by following the co-CRISPR strategy (*28*) using two *nhr-76* crRNAs, *dpy-10* crRNA, tracrRNA, and Cas9 protein. *nhr-76* deletions were detected by PCR and confirmed by Sanger sequencing. Transgenic animals were generated by microinjection following the standard method (*29*). For the transgenic rescue experiments, PCR products of an *nhr-76* genomic fragment at 5 ng/µl or *nhr-76* cDNA under the tissue/cell-specific promoters at 25 ng/µl were injected into NUJ571 *nhr-76(knj51*). For the overexpression experiment, *nhr-76* cDNA under the AWA-specific promoter at 50 ng/µl was injected into N2. For the behavioral assays, co-injection markers (*rps-0*p::HygR at 10 ng/µl and coelomocyte RFP at 25 ng/µl) were injected together with each plasmid. Transgenic animals carrying extrachromosomal arrays were selected by the resistance to 0.125 mg/ml hygromycin in the NGM plate and confirmed by the red fluorescence in coelomocytes. At least two lines were tested. For confocal imaging, tagRFP under the AWA-specific promoter at 25 ng/µl was injected into N2 with a co-injection marker (pRF4 *rol-6(su1006*) at 25 ng/µl). pUC19 plasmid was used to adjust the total concentration of the injected DNA to 100 ng/µl.

### Diacetyl quantification

Diacetyl quantification was performed as previously described (*30*). *L. paracasei* was precultured with MRS (BD Difco™ *Lactobacilli* MRS Broth), minimal MRS (5.0 g Yeast extract, 0.1 g MgSO_4_·7H_2_O, 0.07 g MnSO_4_·5H_2_O, 2.0 g NaH_2_PO_4_, 1.0 g Tween-80 in 1 l of Milli-Q water, pH was adjusted to 5.5), Glucose and citrate-supplemented (6.3 g Glucose and 2.5 g Trisodium citrate dihydrate) minimal MRS, or Glucose and pyruvate-supplemented (6.7 g Glucose and 2.12 g Sodium pyruvate) minimal MRS (*10*). Precultured *L. paracasei* were seeded on 1.5% agar in respective media and incubated for 3 days at 37°C. Two solutions, 325 µl of saturated creatin (0.2 g of creatine into 10 ml of Milli-Q water) and 150 µl of Solution B (3%(w/v) NaOH, 3.5%(w/v) α-naphthol), were mixed in a 1.5 ml tube right before the assay. The same size of agar plugs with and without bacteria were cut out with a P1000 tip, added to the solution above, and crushed well with a P1000 tip. After mixing with a vortex mixer for 15 seconds, the solution was centrifuged at 10,000 rpm for 1 minute at room temperature and filtrated with 0.22 µm MILLEX® GV filter (0.22 µm, Millipore) to remove the agar. The solution was left at room temperature for 30 minutes, and 525-nm absorbance was measured using the DS-11 spectrophotometer (DeNovix).

### Population food-taxis assay

The population food-taxis assay was based on the population chemotaxis assay described above, except that bacterial agar plugs instead of chemicals were used as an attractant. *L. paracasei* was prepared using the same method as diacetyl quantification, using agar plates with pyruvate-supplemented minimal MRS. Cultured plates with bacteria were equilibrated at room temperature in a box before the food-taxis assay. Agar plugs with and without bacteria were cut out with a P1000 tip and placed on the lid of the attractant and control side of an assay plate, respectively.

### Brood size measurement

P_0_ L4 animals were transferred to individual plates 48 hours after egg preparation and transferred to a new plate daily until they stopped depositing eggs. The number of F_1_ progeny was counted under a dissection microscope when they became adults.

### *C. elegans* growth

*E. coli* OP50 and *L. paracasei* were cultured for 16 hours at 37°C with LB broth and MRS, respectively. The cultured bacteria were centrifuged at 7,000 rpm for 10 minutes at 4°C. The pellets were resuspended with 0.9% NaCl and centrifuged at 7,000 rpm for 10 minutes at 4°C. Resuspending and centrifugation were repeated once more, and the supernatants were removed. The pellets were diluted with Nematode Growth buffer (NG buffer, 3 g NaCl, 1 ml 1 M CaCl_2_, 1 ml 1 M MgSO_4,_ and 25 ml 1 M K-PO_4_ in 1 l of Milli-Q water) at 100 mg/ml. Two hundred µl of 100 mg/ml bacteria were seeded on peptone-free NGM plates. In the mixed bacterial food condition, *E. coli* and *L. paracasei* were mixed at the same ratio immediately before seeding. The seeded plates were dried for 24 hours. Synchronized eggs were placed on the bacterial plates, and the growth of animals was observed 72 hours later at 23°C. The images were taken using the same method as the locomotion assay.

### Statistical analysis

Graphs were generated by using R (*31*). Statistical analysis was done with EZR (*32*). Behavioral assay data sets were analyzed with a non-parametric test (Mann-Whitney U test for a single comparison and Kruskal-Wallis test for multiple comparisons). Parametric tests (Student’s t-test and one-way ANOVA) were used for other data sets. In box and whisker plots, the box, vertical line, and whiskers indicate the first and third quartiles, the median, and the max and minimum values excluding outliers, respectively. In bar plots, the standard error of the mean is indicated as a black line.

## Acknowledgments

We thank Noma group members for their invaluable discussion and comments on this manuscript. Some strains were provided by Dr. Shohei Mitani of the National Bioresource Project of Japan and the CGC, which is funded by NIH Office of Research Infrastructure Programs (P40 OD010440).

## Author contributions

Conceptualization: KN

Methodology: RY, KN

Investigation: RY, KN

Visualization: RY

Funding acquisition: KN

Project administration: KN

Supervision: KN

Writing – original draft: RY, KN

Writing – review & editing: RY, KN

## Data and materials availability

All data are available in the main text or the supplementary materials.

## References

1. C. Lopez-Otin, M. A. Blasco, L. Partridge, M. Serrano, G. Kroemer, Hallmarks of aging: An expanding universe. Cell 186, 243–278 (2023).

2. C. I. Bargmann, E. Hartwieg, H. R. Horvitz, Odorant-selective genes and neurons mediate olfaction in C. elegans. Cell 74, 515–527 (1993).

3. J. I. Choi, K. H. Yoon, S. Subbammal Kalichamy, S. S. Yoon, J. Il Lee, A natural odor attraction between lactic acid bacteria and the nematode Caenorhabditis elegans. ISME J 10, 558–567 (2016).

4. P. Sengupta, J. H. Chou, C. I. Bargmann, odr-10 encodes a seven transmembrane domain olfactory receptor required for responses to the odorant diacetyl. Cell 84, 899–909 (1996).

5. N. Suryawinata et al., Dietary E. coli promotes age-dependent chemotaxis decline in C. elegans. Scientific Reports 14, (2024).

6. S. Ward, J. S. Carrel, Fertilization and sperm competition in the nematode Caenorhabditis elegans. Dev Biol 73, 304–321 (1979).

7. T. Noble, J. Stieglitz, S. Srinivasan, An integrated serotonin and octopamine neuronal circuit directs the release of an endocrine signal to control C. elegans body fat. Cell Metab 18, 672–684 (2013).

8. M. E. Colosimo, S. Tran, P. Sengupta, The divergent orphan nuclear receptor ODR-7 regulates olfactory neuron gene expression via multiple mechanisms in Caenorhabditis elegans. Genetics 165, 1779–1791 (2003).

9. P. Sengupta, H. A. Colbert, C. I. Bargmann, The C. elegans gene odr-7 encodes an olfactory-specific member of the nuclear receptor superfamily. Cell 79, 971–980 (1994).

10. B. D. Jyoti, A. K. Suresh, K. V. Venkatesh, Diacetyl production and growth of Lactobacillus rhamnosus on multiple substrates. World Journal of Microbiology and Biotechnology 19, 509–514 (2003).

11. K. Kawamura, I. N. Maruyama, Forward Genetic Screen for Caenorhabditis elegans Mutants with a Shortened Locomotor Healthspan. G3 (Bethesda) 9, 2415–2423 (2019).

12. D. F. Midkiff, J. Huayta, J. D. Lichty, J. P. Crapster, A. San-Miguel, Identifying C. elegans lifespan mutants by screening for early-onset protein aggregation. iScience 25, 105460 (2022).

13. S. N. Austad, J. M. Hoffman, Is antagonistic pleiotropy ubiquitous in aging biology? Evol Med Public Health 2018, 287–294 (2018).

14. G. C. Williams, PLEIOTROPY, NATURAL SELECTION, AND THE EVOLUTION OF SENESCENCE. Evolution 11, 398–411 (1957).

15. H. G. Son, O. Altintas, E. J. E. Kim, S. Kwon, S. V. Lee, Age-dependent changes and biomarkers of aging in Caenorhabditis elegans. Aging Cell 18, e12853 (2019).

16. E. Spanoudakis, N. Tavernarakis, Age-associated anatomical and physiological alterations in Caenorhabditis elegans. Mech Ageing Dev 213, 111827 (2023).

17. J. J. Collins, C. Huang, S. Hughes, K. Kornfeld, The measurement and analysis of agerelated changes in Caenorhabditis elegans. WormBook, 1–21 (2008).

18. A. a. P. A. Aranda, Nuclear Hormone Receptors and Gene Expression. Physiological Reviews 81, 1269–1304 (2001).

19. B. G. Fenyves et al., Dual Role of an mps-2/KCNE-Dependent Pathway in Long-Term Memory and Age-Dependent Memory Decline. Curr Biol 31, 527–539 e527 (2021).

20. B. S. Samuel, H. Rowedder, C. Braendle, M.-A. Félix, G. Ruvkun, Caenorhabditis elegans responses to bacteria from its natural habitats. Proceedings of the National Academy of Sciences 113, E3941-E3949 (2016).

21. B. M. Aleogho et al., Aberrant neuronal hyperactivation causes an age- and diet-dependent decline in associative learning behavior. bioRxiv, 2024.2003.2021.586045 (2024).

22. S. Brenner, The genetics of Caenorhabditis elegans. Genetics 77, 71–94 (1974).

23. T. Stiernagle, Maintenance of C. elegans. (WormBook, 2006).

24. C. A. Schneider, W. S. Rasband, K. W. Eliceiri, NIH Image to ImageJ: 25 years of image analysis. Nature methods 9, 671–675 (2012).

25. F. G. Mann, E. L. Van Nostrand, A. E. Friedland, X. Liu, S. K. Kim, Deactivation of the GATA Transcription Factor ELT-2 Is a Major Driver of Normal Aging in C. elegans. PLOS Genetics 12, e1005956 (2016).

26. S. K. Han et al., OASIS 2: online application for survival analysis 2 with features for the analysis of maximal lifespan and healthspan in aging research. Oncotarget 7, 56147–56152 (2016).

27. D. G. Gibson et al., Enzymatic assembly of DNA molecules up to several hundred kilobases. Nature methods 6, 343–345 (2009).

28. H. Kim et al., A co-CRISPR strategy for efficient genome editing in Caenorhabditis elegans. Genetics 197, 1069–1080 (2014).

29. C. C. Mello, J. M. Kramer, D. Stinchcomb, V. Ambros, Efficient gene transfer in C. elegans: extrachromosomal maintenance and integration of transforming sequences. EMBO J 10, 3959–3970 (1991).

30. J. Mattessich, J. R. Cooper, The spectrophotometric determination of diacetyl.

31. R_Core_Team, R: A language and environment for statistical computing. R Foundation for Statistical Computing. (2020).

32. Y. Kanda, Investigation of the freely available easy-to-use software ‘EZR’ for medical statistics. Bone Marrow Transplantation 48, 452–458 (2013).

33. Q. Ch’Ng et al., Identification of Genes That Regulate a Left-Right Asymmetric Neuronal Migration in Caenorhabditis elegans. Genetics 164, 1355–1367 (2003).

