## Supplemental data for "A nuclear hormone receptor *nhr-76* programs age-dependent chemotaxis decline in *C. elegans*"

#### Supplementary Text

**Figure S1. Transient FUdR treatment and mating increase the number of progeny in aged animals.** The total number of progeny produced by Day 3, 4, and 5 animals was counted. For the FUdR condition, animals were treated with FUdR for 24 hours from L4 (48 hours after epp preparation). For the condition with males, an excess number of young males were added to Day 3, 4, or 5 hermaphrodites. Statistical tests were conducted using One-way ANOVA with Tukey's test. ns: p>0.05; **p<0.01; ***p<0.001.

**Figure S2. Aged animals showed locomotion defects.** The locomotion speed of Day1 to Day15 animals was measured on food for wt and *knj39* mutants. *knj39* mutants showed wild-type-like defect of locomotion speed on Day15. Statistical tests were conducted using One-way ANOVA with the Dunnett's test. *p<0.05; **p<0.01; ***p<0.001.

**Figure S3. *nhr-76(knj39)* is semidominant.** After crossing wild-type (+/+) or *nhr-76(knj39)* animals with males carrying the GFP marker (gfp, *muIs32[mec-7p::GFP]*), GFP-positive $F_1$ animals were counted as the heterozygotes in chemotaxis assays. Statistical tests were conducted using the Kruskal-Wallis with Steel-Dwass test. ns: p>0.05; *p<0.05.

**Figure S4. *nhr-76* mutants do not show high chemotaxis ability on Day1.** The chemotaxis assay was conducted with 0.01% diacetyl for *nhr-76(knj39)* on Day1. The statistical test was conducted using the Mann-Whitney U test. ns: p>0.05.

**Figure S5. *nhr-76(knj51)* mutants have a normal reproductive span.** (A) and (C) The total number of progeny was not significantly different between wt and *nhr-76(knj51)* mutants. (B) and (D) The reproductive span of the wild-type, *nhr-76(knj39)* and *nhr-76(knj51)* mutants. The number of progeny deposited during the indicated period was normalized with the total number of progeny of each genotype. Statistical tests were conducted using Student's t-test for (**A and C**). ns: p>0.05; **p<0.01.

**Figure S6. Aged *knj39* mutants require *odr-10* for diacetyl chemotaxis.** The chemotaxis assays were conducted with 0.1% diacetyl for the wild type, *nhr-76(knj39), odr-10(ky225),* and *nhr-76(knj39); odr-10(ky225)* on Day1 and Day5. Statistical tests were conducted using Kruskal-Wallis with Steel tests against *odr-10; nhr-76* double mutant counterparts. ns: p>0.05; ***p<0.01. The data of the wild type and *knj39* mutants are the same as in Figure 3A.

**Figure S7. Tissue/cell-specific rescue experiment for *nhr-76* mutants.** Chemotaxis assays in Figure 2C were conducted with transgenic lines with different extrachromosomal arrays. Statistical tests were conducted using Kruskal-Wallis with Steel test against *nhr-76(knj51)* mutant counterparts. ns: p>0.05; **p<0.01. The data of the wild type is the same as in Figure 2C.

**Figure S8. Mutants defective in lipid metabolisms do not ameliorate age-dependent chemotaxis decline.** Chemotaxis assays toward 0.1% diacetyl chemotaxis were conducted with indicated genotypes and ages. Statistical tests were conducted using Kruskal-Wallis with Steel test against wild-type counterparts. ns: p>0.05; *p<0.05; **p<0.01; ***p<0.001.

**Figure S9. *nhr-76* mutants do not ameliorate age-dependent chemotaxis decline toward benzaldehyde.** Chemotaxis assays of the wild-type and *nhr-76(knj39)* were conducted with

0.01% benzaldehyde, sensed by AWC sensory neurons. Statistical tests were conducted using the Kruskal-Wallis with Steel-Dwass test. ns: $p>0.05$; **$p<0.01$.

**Figure S10. *nhr-76* overexpression does not decrease the chemotaxis ability on Day1.** Chemotaxis assays in Figure 3F were conducted with transgenic lines with different extrachromosomal arrays. Statistical tests were conducted using the Kruskal-Wallis with Steel-Dwass test. Different alphabets indicate significant differences.

**Figure S11. *L. paracasei* does not support the growth of *C. elegans* larvae.** Representative images showing the animals cultured for 3 days from eggs with *E. coli*, mixed bacteria of *E. coli* and *L. paracasei*, or *L. paracasei*. Scale bar: 1 mm.

### Figure S1 Yokosawa and Noma

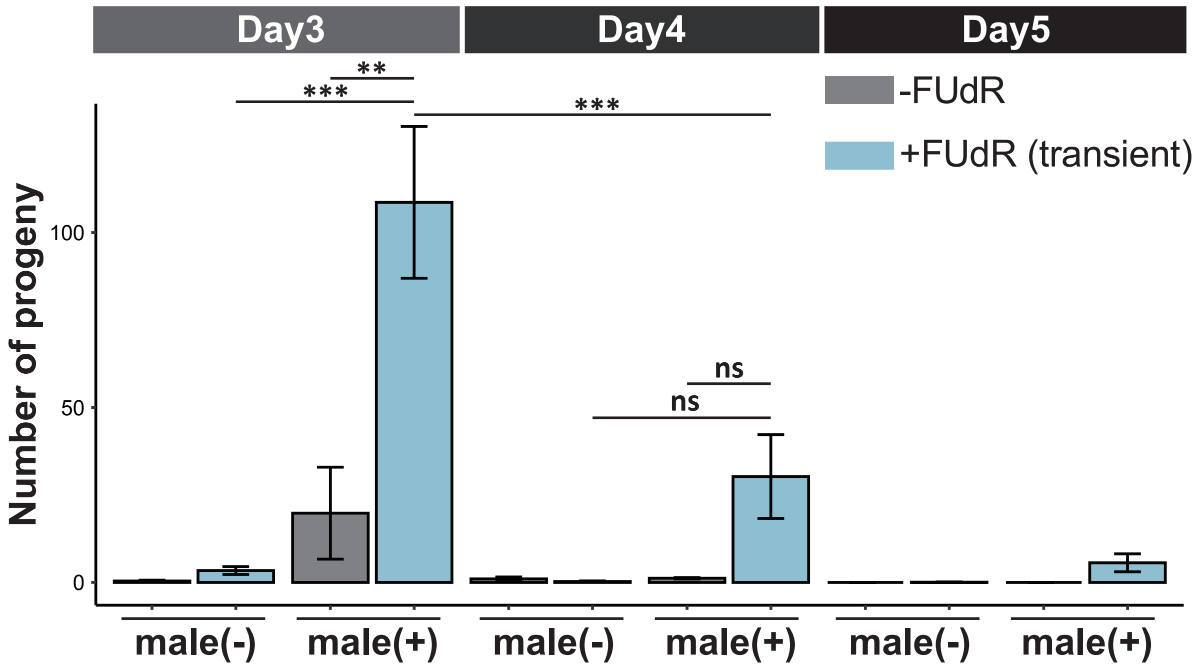

### Figure S2

Yokosawa and Noma

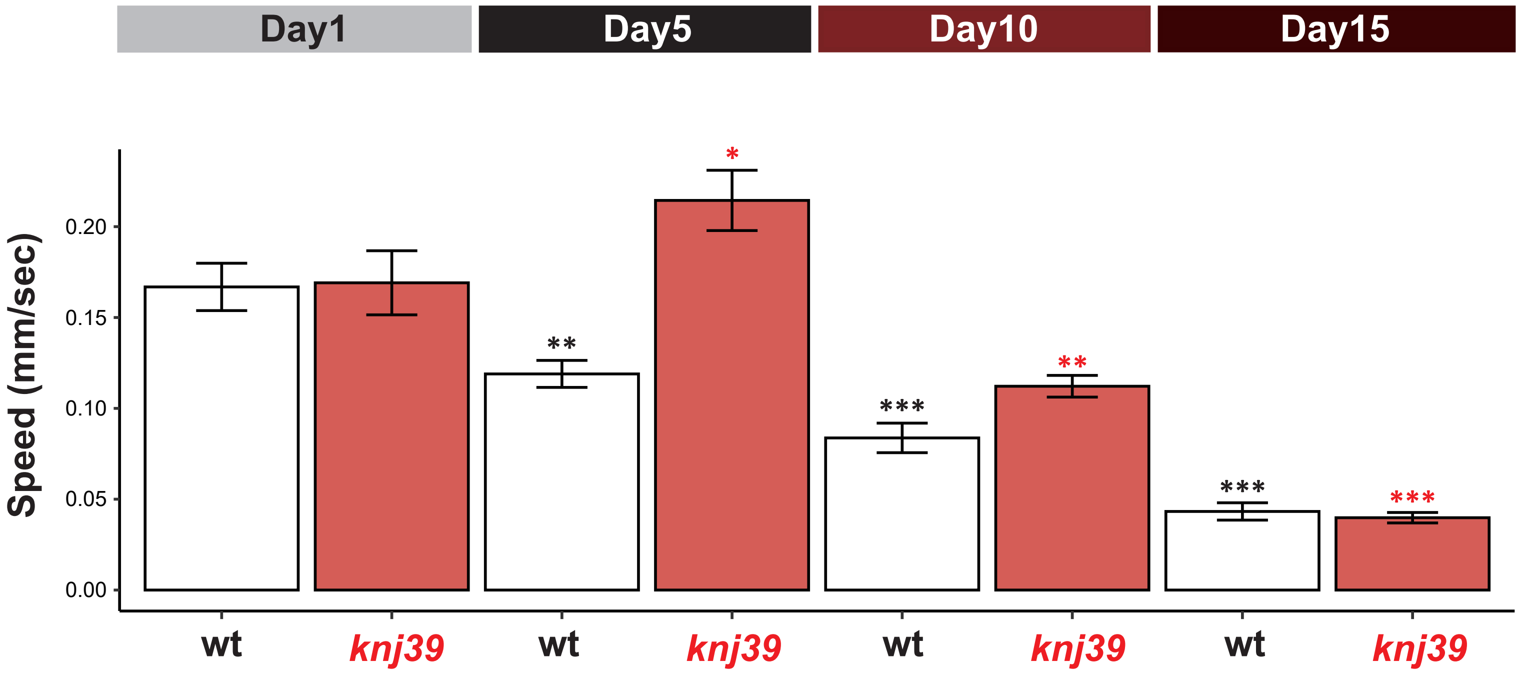

### Figure S3

Yokosawa and Noma

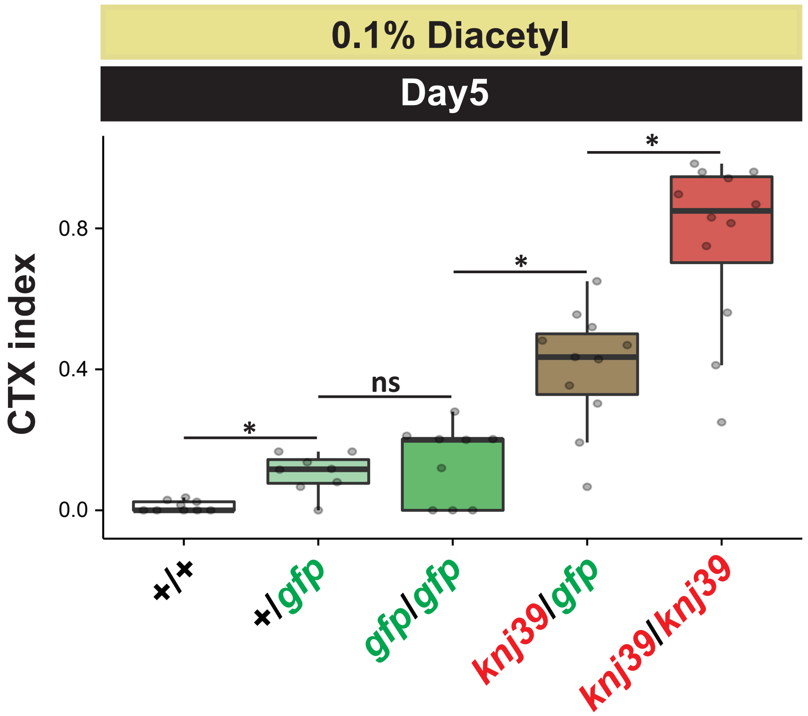

### Figure S4

Yokosawa and Noma

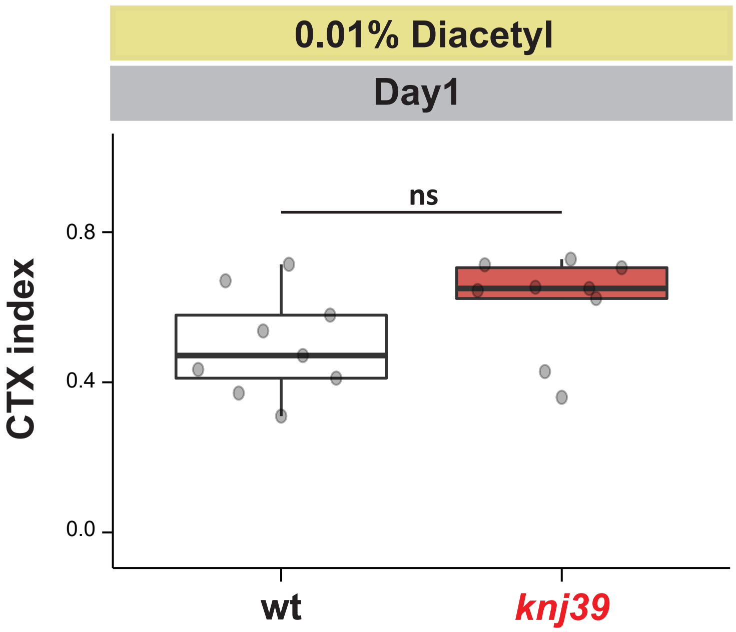

### Figure S5

Yokosawa and Noma

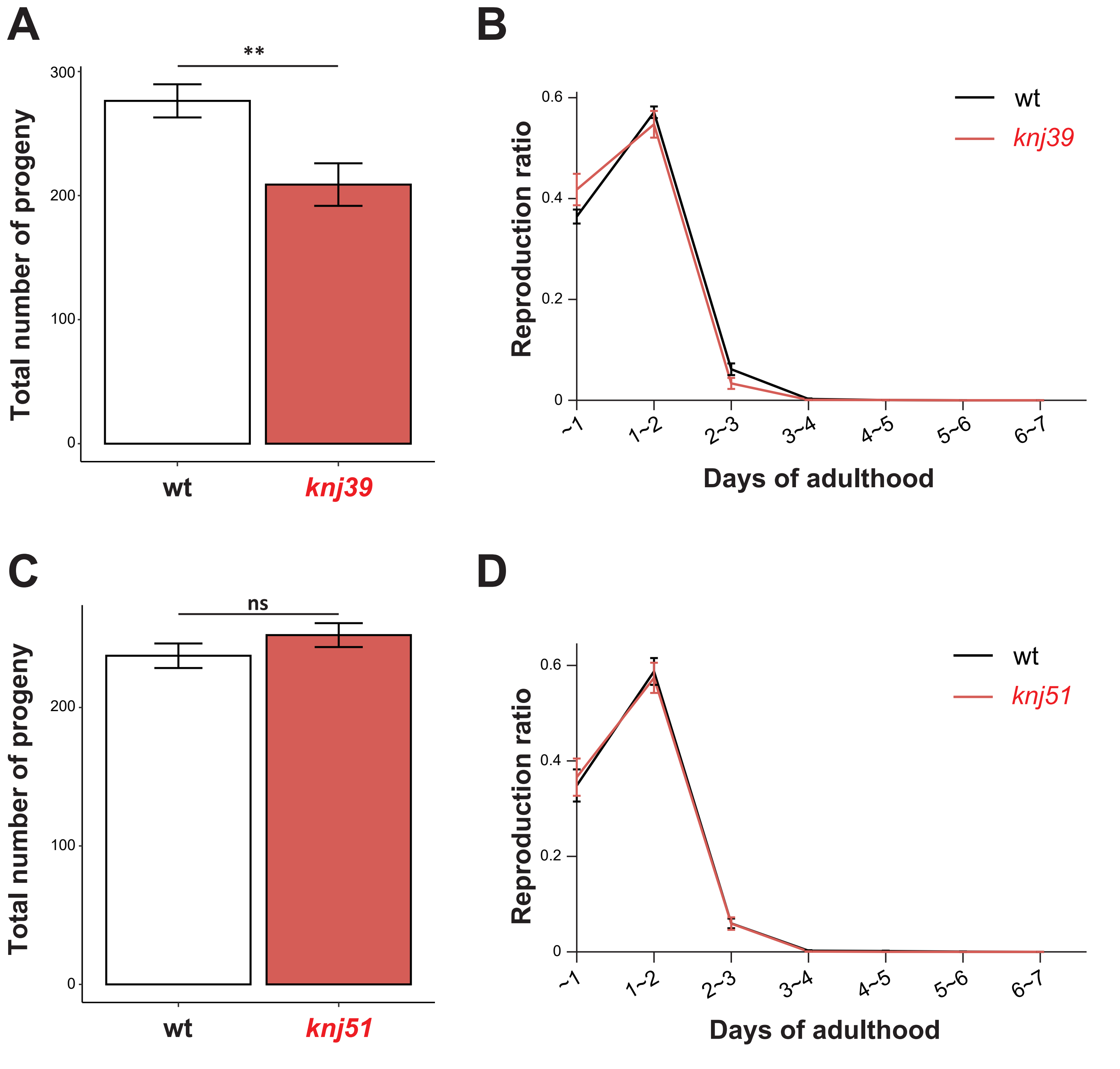

### Figure S6

### Yokosawa and Noma

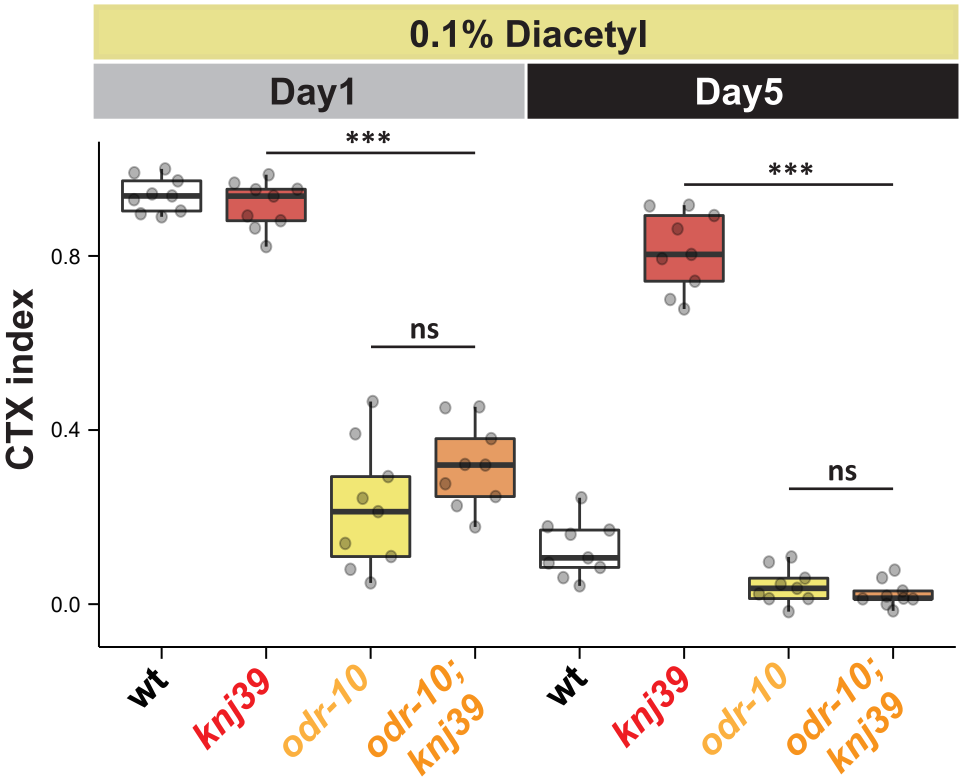

### Figure S7

Yokosawa and Noma

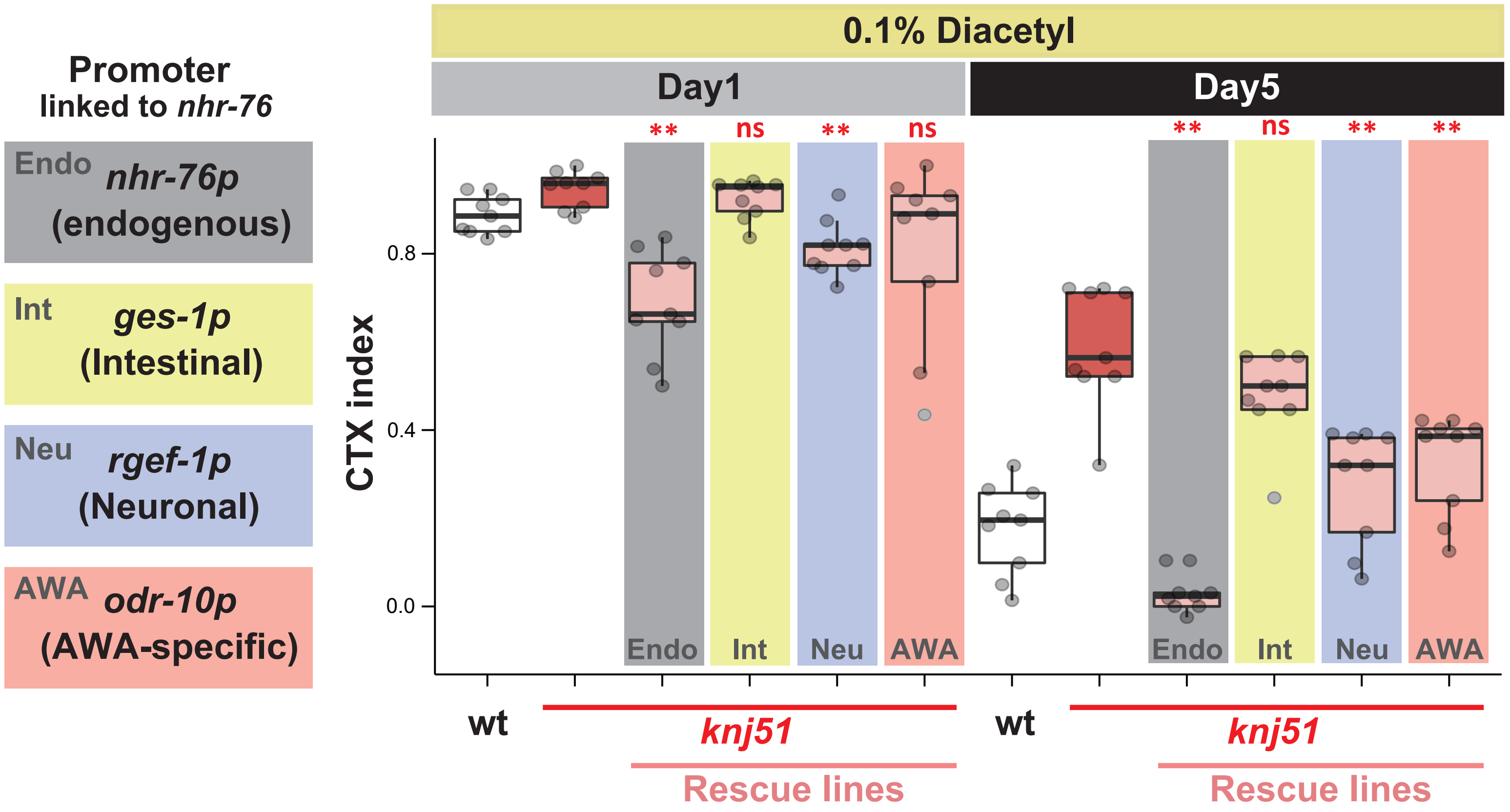

### Figure S8

Yokosawa and Noma

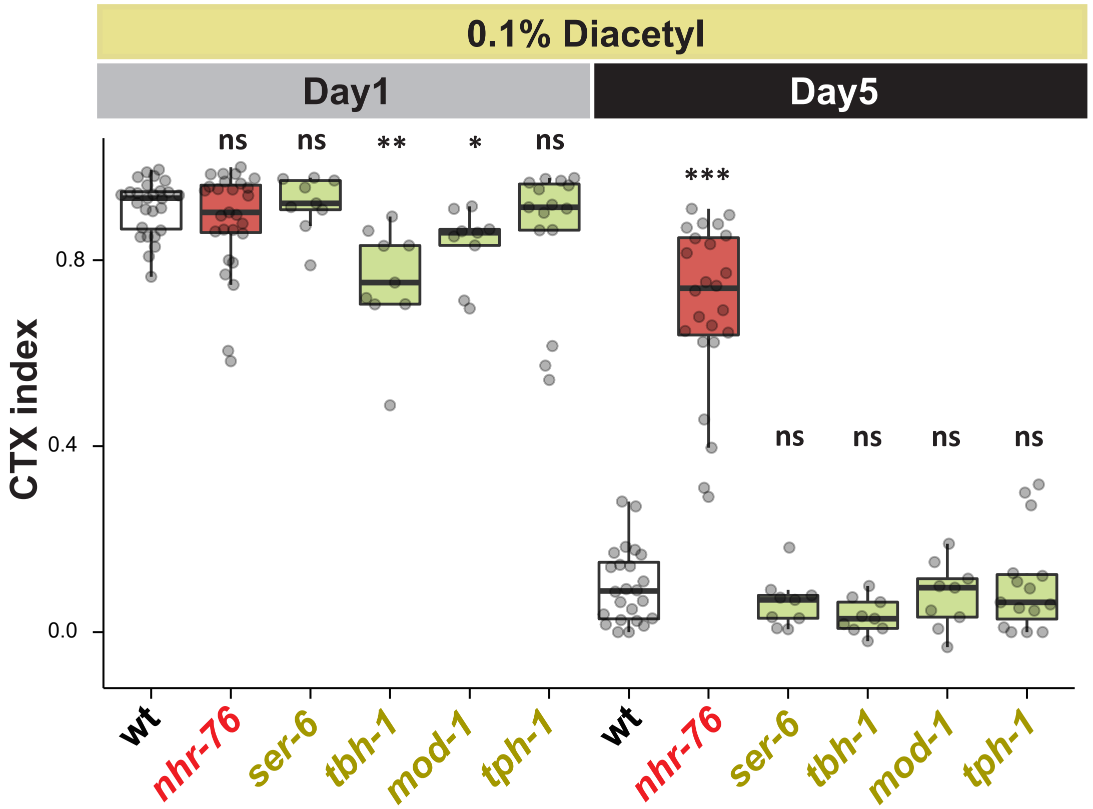

### Figure S9

Yokosawa and Noma

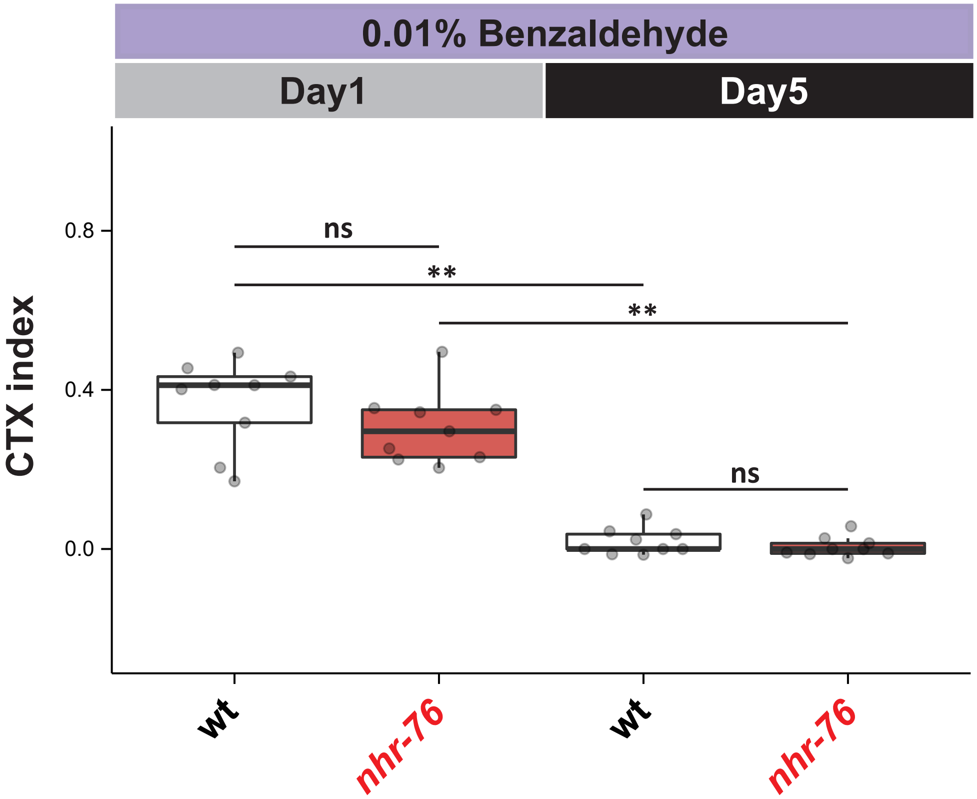

### Figure S10

Yokosawa and Noma

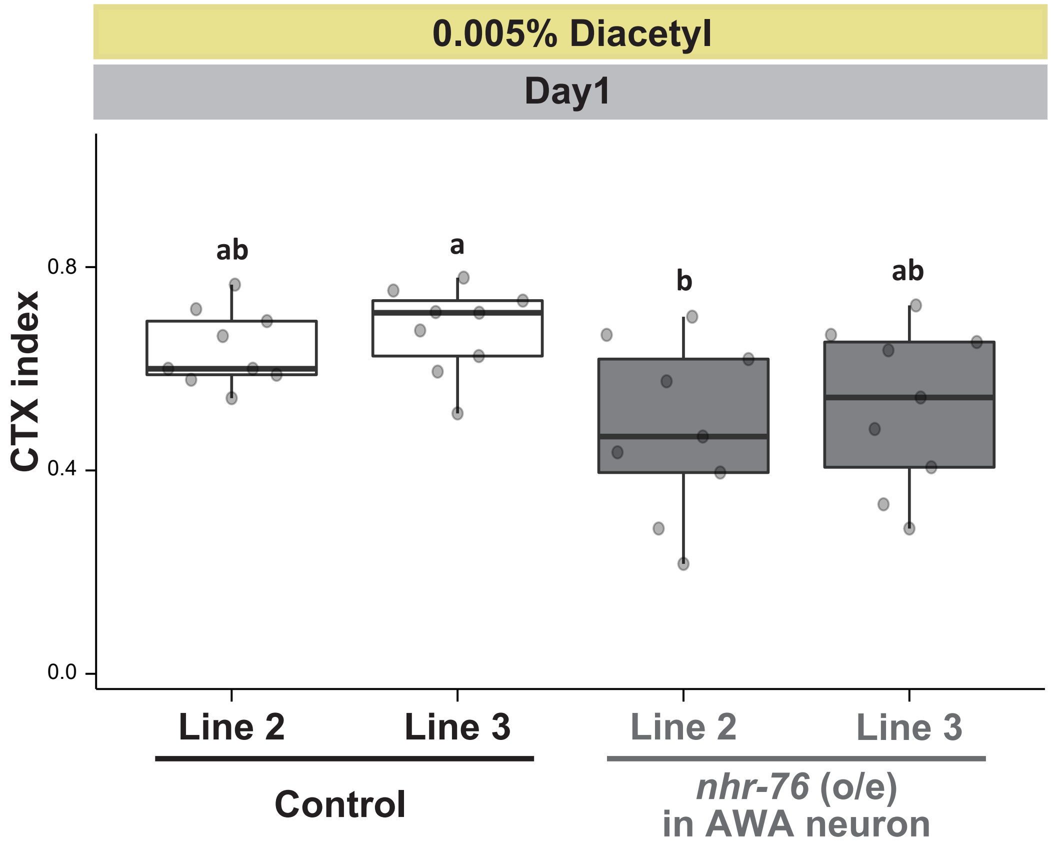

Figure S11 Yokosawa and Noma

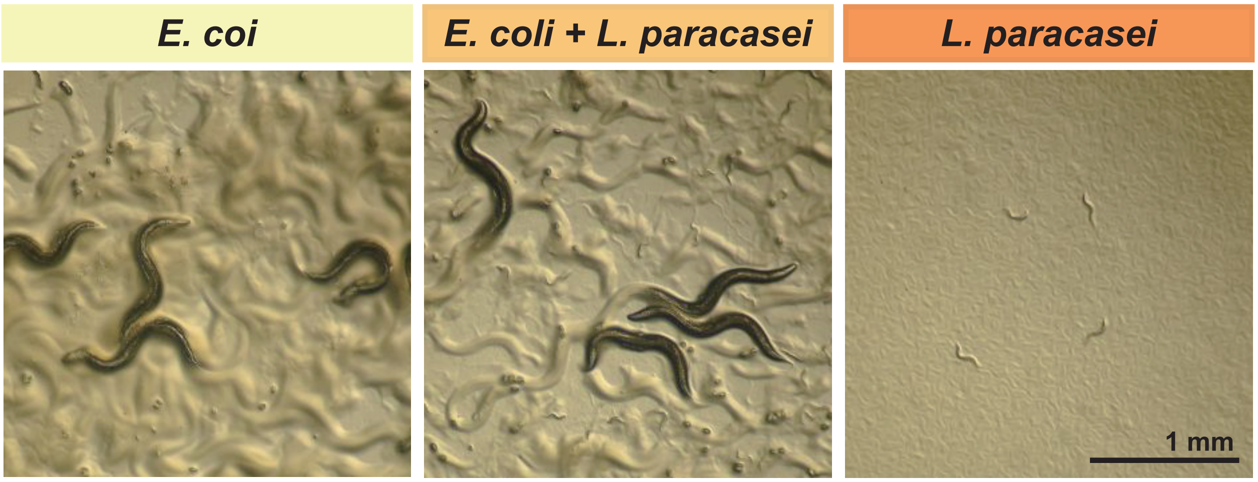

**Table S1** ***C. elegans*** **strains**

| Strain | Genotype | Note | Source | Figure |
|---|---|---|---|---|
| N2 | *wild type* | | CGC | 1B, 1C, 1E, 2B, 2C, 3A, 3B, 3C, 3D, 4C, S1, S2, S3, S5A, S5B,S5, S5C, S5D, S6, S7, S8, S9, S11 |
| NUJ392 | *nhr-76(knj39) IV* | 2x outcrossed *knj39* | This study, from forward genetic screen | S3, S4 |
| NUJ455 | *nhr-76(knj39) IV* | 3x outcrossed *knj39* | This study, Cross | 3C, 3D, S3, S9 |
| NUJ553 | *nhr-76(knj39) IV* | 4x outcrossed *knj39* | This study, Cross | 1B, 1E, 2B, S2, S8 |
| NUJ557 | *nhr-76(knj39) IV* | 5x outcrossed *knj39* | This study, Cross | 1B, 1C, 1E, 2B, 3A, 3B, 3C, 3D, 4C, S2, S5A, S5B, S6, S9 |
| NUJ580 | *kyIs53[odr-10::GFP] X* | 4x outcrossed strain of CX3344 *kyIs53[odr-10::GFP] X* | This study, Cross | 1D |
| NUJ648 | *nhr-76(knj39) IV; kyIs53[odr-10::GFP] X* | | This study, Cross | 1D |
| NUJ560 | *nhr-76(tm671) IV* | 2x outcrossed strain of *nhr-76(tm671)* from NBRP | This study, Cross | 2B |
| NUJ571 | *nhr-76(knj51) IV* | This study, injection | This study, CRISPR | 2B, S5C, S5D |
| NUJ603 | *nhr-76(knj52) IV* | This study, injection | This study, CRISPR | 2B |
| NUJ588 | *nhr-76(knj51) IV; knjEx235[ccRFP + rps-0p::HygR]* | Control for the co-injection markers (Hygromicine resistantce and Coelomocyte RFP) | This study, Injection | 2C |
| NUJ576 | *nhr-76(knj51) IV; knjEx226[nhr-76(gDNA) + ccRFP + rps-0p::HygR]* | Rescue of *nhr-76* with a PCR-amplified genomic fragment | This study, Injection and Cross | 2C |
| NUJ607 | *nhr-76(knj51) IV; knjEx248[ges-1p::nhr-76(cDNA) + ccRFP + rps-0p::HygR]* | Intestinal rescue of *nhr-76* | This study, Injection | 2C |
| NUJ605 | *nhr-76(knj51) IV; knjEx246[rgef-1p::nhr-76(cDNA) + ccRFP + rps-0p::HygR]* | Pan-neuronal rescue of *nhr-76* | This study, Injection | 2C |
| NUJ609 | *nhr-76(knj51) IV; knjEx250[odr-10p::nhr-76(cDNA) + ccRFP + rps-0p::HygR]* | AWA-specific rescue of *nhr-76* | This study, Injection | 2C |
| CX4 | *odr-7(ky4) X* | | CGC | 3A, 3B |
| NUJ624 | *nhr-76(knj39) IV; odr-7(ky4) X* | | This study, Cross | 3A, 3B |
| NUJ636 | *wgIs203[nhr-76::TY1::EGFP::3xFLAG + unc-119(+)]; knjEx93[odr-10p::tagRFP + rol-6(su1006)]* | *knjEx93* was crossed with OP203 *unc-119 (tm4063) III; wgIs203 [nhr-76::TY1::EGFP::3xFLAG + unc-119(+)]* from CGC. It may contain *unc-119(tm4063)* . | This study, Cross | 3E |
| NUJ627 | *knjEx234[ccRFP + rps-0p::HygR]* | Co-injection marker control (Hygromicine resistantce and Coelomocyte RFP) (Line1) | This study, Cross | 3F |
| NUJ634 | *knjEx261[odr-7p::nhr-76(cDNA) + ccRFP + rps-0p::HygR]* | AWA-specific *nhr-76* over expression (Line1) | This study, Injection | 3F |
| CZ10969 | *muIs32[mec-7p::GFP + lin-15(+)] II* | Used for ganerating heterozygous animals in dominancy test | Ref. (*33* ) | S3 |
| NUJ457 | *odr-10(ky225) X* | 1x outcrossed strain of CX3410 *odr-10(ky225)X* from CGC | This study, Cross | S6 |
| NUJ623 | *nhr-76(knj39) IV; odr-10(ky225) X* | | This stury, Cross | S6 |
| NUJ587 | *nhr-76(knj51) IV; knjEx234[ccRFP + rps-0p::HygR]* | Co-injection marker control (Hygromicine resistantce and Coelomocyte RFP) (Line 1) | This study, Injection | S7 |
| NUJ590 | *nhr-76(knj51) IV; knjEx227[hr-76(gDNA) + ccRFP + rps-0p::HygR]* | Genomic fragment rescue of *nhr-76* | This study, Injection and Cross | S7 |
| NUJ608 | *nhr-76(knj51) IV; knjEx249[ges-1p::nhr-76(cDNA) + ccRFP + rps-0p::HygR]* | Intestin-specific rescue of *nhr-76* | This study, Injection | S7 |
| NUJ612 | *nhr-76(knj51) IV; knjEx253[rgef-1p::nhr-76(cDNA) + ccRFP + rps-0p::HygR]* | Pan-neuronal rescue of *nhr-76* | This study, Injection | S7 |
| NUJ611 | *nhr-76(knj51) IV; knjEx252[odr-10p::nhr-76(cDNA) + ccRFP + rps-0p::HygR]* | AWA-specific rescue of *nhr-76* | This study, Injection | S7 |
| IK1348 | *ser-6(tm2146) IV* | 2x outcrossed *ser-6(tm2146)* from NBRP | A gift from Ikue Mori | S8 |
| MT9455 | *tbh-1(n3247) X* | | CGC | S8 |
| MT9668 | *mod-1(ok103) V* | | CGC | S8 |
| NUJ578 | *tph-1(mg280) II* | 1x backcrossed strain of MT15434 *tph-1(mg280) II* from CGC | This study, Cross | S8 |
| NUJ628 | *knjEx235[ccRFP + rps-0p::HygR]* | Co-injection marker control (Hygromicine resistantce and Coelomocyte RFP) (Line 2) | This study, Injection and Cross | S10 |
| NUJ629 | *knjEx236[ccRFP + rps-0p::HygR]* | Co-injection marker control (Hygromicine resistantce and Coelomocyte RFP) (Line 3) | This study, Injection and Cross | S10 |
| NUJ633 | *knjEx260[odr-7p::nhr-76(cDNA) + ccRFP + rps-0p::HygR]* | AWA-specific *nhr-76* over expression (Line2) | This study, Injection | S10 |
| NUJ635 | *knjEx262[odr-7p::nhr-76(cDNA) + ccRFP + rps-0p::HygR]* | AWA-specific *nhr-76* over expression (Line3) | This study, Injection | S10 |

**Table S2 Primers and crRNAs**

| Primer | Gene | Sequence (5' to 3') | Note |
|---|---|---|---|
| KN1170 | *cdc-42* | CTGCTGGACAGGAAGATTACG | For qPCR (as a reference) |
| KN1171 | *cdc-42* | CTCGGACATTCTCGAATGAAG | For qPCR (as a reference) |
| KN1624 | *odr-10* | TTAGTACATTGGTGACAGCCGC | For qPCR |
| KN1625 | *odr-10* | TTGGAATCGGCGCCAGACGG | For qPCR |
| KN1699 | *odr-7* | CAAGCCGCGATGGAACTGGG | For qPCR |
| KN1700 | *odr-7* | GTGCTGCACATTTTCTGGAAGC | For qPCR |
| KN2076 | *nhr-76* | GTCGGCTATCTCTGGTGACG | For generating genomic fragment of *nhr-76* |
| KN2077 | *nhr-76* | GCCCACTACTTAGACCACGAG | For generating genomic fragment of *nhr-76* |
| KN2087 | *nhr-76* | TTTGTACAAAAAAGCAGGCTCCGAATTCATGGAGGTGCTCGGGAAGCAG | For generating *nhr-76* cDNA |
| KN2088 | *nhr-76* | TTGTACAAGAAAGCTGGGTCGAATTCTTACGTGAACGCGAGATCATCAA | For generating *nhr-76* cDNA |
| KN1251 | *dpy-10* | *GCUACCAUAGGCACCACGAGGUUUUAGAGCUAUGCU* | crRNA for Co-CRISPR strategy |
| KN2082 | *nhr-76* | CUGCCAGUUGGGGAAGUAUGGUUUUAGAGCUAUGCU | crRNA for CRISPR KO |
| KN2083 | *nhr-76* | GCUGAUGGAAGGCACGAGAAGUUUUAGAGCUAUGCU | crRNA for CRISPR KO |

**Table S3 Plasmids**

| Plasmid | Description | Related strains | Note |
|---|---|---|---|
| pCZGY66 | *rgef-1p::GTW-unc-54 3'UTR* | | Gift from Yishi Jin |
| pKEN838 | *ges-1p::GTW-3'UTR(unc-54)* | | |
| pKEN927 | *odr-7p::GTW-3'UTR(unc-54)* | | |
| pKEN928 | *odr-10p::GTW-3'UTR(unc-54)* | | |
| pKEN1071 | *nhr-76(cDNA)* pCR8 Backborne | | |
| pKEN1072 | *rgef-1p::nhr-76(cDNA)* | NUJ605, NUJ612 | pKEN1071 + pCZGY66 |
| pKEN1073 | *ges-1p::nhr-76(cDNA)* | NUJ607, NUJ608 | pKEN1071 + pKEN838 |
| pKEN1074 | *odr-10p::nhr-76(cDNA)* | NUJ609, NUJ611 | pKEN1071 + pKEN927 |
| pKEN1075 | *odr-7p::nhr-76(cDNA)* | NUJ633, NUJ634, NUJ635 | pKEN1071 + pKEN928 |
| pKEN939 | *odr-10p::tagRFP* | NUJ636 | |
| pRF4 | *rol-6(su1006)* | NUJ636 | |
| pUC19 | | NUJ587, NUJ588, NUJ576, NUJ590, NUJ605, NUJ612, NUJ607, NUJ608, NUJ609, NUJ611, NUJ627, NUJ628, NUJ629, NUJ633, NUJ634, NUJ635, NUJ636 | |
| pKEN954 | *rps-0shp::HygR* | NUJ587, NUJ588, NUJ576, NUJ590, NUJ605, NUJ612, NUJ607, NUJ608, NUJ609, NUJ611, NUJ627, NUJ628, NUJ629, NUJ633, NUJ634, NUJ635 | |
| pKEN281 | coelomocyte RFP (ccRFP) | NUJ587, NUJ588, NUJ576, NUJ590, NUJ605, NUJ612, NUJ607, NUJ608, NUJ609, NUJ611, NUJ627, NUJ628, NUJ629, NUJ633, NUJ634, NUJ635 | |
